# Bioinformatics Workflow Management With The Wobidisco Ecosystem

**DOI:** 10.1101/213884

**Authors:** Sebastien Mondet, Bulent Arman Aksoy, Leonid Rozenberg, Isaac Hodes, Jeff Hammerbacher

## Abstract

To conduct our computational experiments, our team developed a set of workflow-management-related projects: Ketrew, Biokepi, and Coclobas. The family of tools and libraries are designed with reliability and flexibility as main guiding principles. We describe the components of the software stack and explain the choices we made. Every piece of software is free and open-source; the umbrella documentation project is available at https://github.com/hammerlab/wobidisco.

## Introduction

### State Of The Union

Over the past half-century the computer and software worlds have proven to be a complete engineering disaster. Extremely poor quality standards, mostly due to humans overestimating their capabilities [1], has lead to the deployment of very unsecure [2] and unreliable [3], [4] software in all sectors of industry and academia. The biomedical research community has added to the phenomenon by allocating very little funding and training to software development; and moreover providing poor recognition for (comparatively) valuable efforts in the domain. Despite some advances in the open-sourcing and community building, like projects using proper online hosting and distribution [5], [6] with decent licenses, and opening access to data, the situation as of 2017 is still that Bioinformatics tools are of extremely poor quality and prove to hurt the productivity of the whole community.

While our projects stay within cancer immunotherapy [7], we aim here at showing solutions for the bioinformatics community at large. Most, if not all, bioinformatics tools presented in various papers, provide partial analysis “steps” blessed through peer-review, and distributed as software packages. Bionformaticians need many of those tools chained together through partially compatible [8] and/or weakly defined file-formats [9], to bring raw data (usually from sequencing platforms) to usable biologically or medically relevant information. We call the whole graph of computations a *pipeline*. Data analysts also need to be able to easily rearrange the ordering of the steps in the pipeline, make their parameters vary, over many similar experiments, while keeping track of the results.

We also want to easily and safely add new tools; adapt to new versions; and install software with as much automation as possible. While we need to assume that anything can randomly fail for obscure reasons, and have to deal with adverse conditions (firewalls, university VPNs, cloud hosting, etc.), we want to optimize infrastructure usage and make reproducible research *easier*. Note that being able to rerun someone else’s analysis bug-for-bug as a “black-box” [10] is very useful but it is not yet proper reproducible research, which should mean being able to re-implement all the tools from their mathematical description/specification, including bugs, imprecisions, and unexpected behaviors if any.

These hurdles on a bioinformatician’s path to publication have lead to a huge demand for pipeline automation tools, a.k.a “Workflow Engines.” A lot of them are hence in use, including many whose development originates from the bioinformatics community itself. We reviewed many of them, cf. Related Work (section), and concluded that none of them came up to our expectations and that their development setup would make it very hard for us to reach an acceptable value by simply contributing. We found that they were not flexible enough, often specific to a platform (e.g. Hadoop), and driven by lots of strong assumptions (e.g. on networks, file-systems, or regarding the topology of the pipeline graphs, etc.). Moreover, we observed very little support for fault-tolerance or guarantees against hard-to-track mistakes. We also argue that the use of custom Domain Specific Languages (DSLs) almost never renders enough flexibility and reliability. Indeed, programmers think they need a very simple DSL, then they realize they need a form of variable-definition-and-substitution, then a “conditional” construct appears, and the list of programming-language-features keeps growing; cf. the *“yet another half-baked Turing-complete language”* syndrome [11], [12].

### Description of The Present Work

To conduct our own computational experiments, we started our own family of workflow-management-related projects: Ketrew, Biokepi, and Coclobas.

- Ketrew is a general-purpose workflow engine (which has also been used for non-bioinformatics applications); it is based on an *Embedded* Domain Specific Language (EDSL), that allowed us to build properly modular libraries of “workflow-steps.”
- Biokepi is our main library collecting and wrapping bioinformatics software tools as assembly-ready pieces of Ketrew pipeline.
- Ketrew has various backend-plugins defining how to communicate with batch scheduling systems; Coclobas is such a new backend; it allows us to harness elastic and container-based infrastructure, such as Kubernetes [13] clusters as provided by the Google Container Engine (GKE, [14]) or AWS-Batch clusters provided by Amazon [15].

This modular setup allowed us to experiment with the *different levels of abstraction* for concisely expressing bioinformatics pipelines. We can see this in our “flagship” application: Epidisco a family of pipelines for “personalized cancer epitope discovery and peptide vaccine prediction” used among other things in the NCT02721043 clinical trial [16], [17]. We created the “umbrella documentation project” named Wobidisco as a centralized entry-point to the family of projects presented in this paper.

This work has been guided by the following general design ideas:

- We use *Embedded* DSLs with static, as strong as possible, type checking; this gives us lot of flexibility to develop, inspect, and maintain very complex pipelines while providing safety guarantees, proper semantics, and always up-to-date “IDE-like” documentation. In particular, Biokepi’s high-level pipeline EDSL (section), is based on recent research by Suzuki et al. [18] which provides flexible and well-typed extensibility.
- Small well-defined abstractions work better than monumental piles of spaghetti. This *modularity* is also important for manageable documentation efforts.
- We acknowledge that we cannot handle or even envision all possible use-cases; every (layer of) abstraction has to be extensible and “escapable” by the users.
- We aim at stressing correctness and fault-tolerance.
- Since system-administration (or “DevOps”) are often missing or under-staffed/underfunded; we need to make deployment in adverse conditions as easy and flexible as possible.
- We want to maximize open/free-for-anything availability of software and data [19].

Many of the above guide-lines lead us to use a *saner-than-usual* implementation language: OCaml [20]. The above choices may not be “traditional,” we discuss them in the following section.

### Quick Digression: Types (and OCaml)

Within bioinformatics tools as with most low quality general purpose software, we observe a lot of overestimation of programmers’ abilities to produce well designed and carefully implemented programs. Indeed, most dismiss or misunderstand the importance of *types*, we see this through the use of popular but *unityped* (as in “one type”) or unsound languages like Python, Perl, Ruby, or C++.

Type theory is an expression of constructive logic which is, or should be, to software engineering what calculus and physics are to civil engineering. Types are not only a great way of thinking about (and modeling) programming problems; types are useful logical properties describing (pieces of) a program. Checking types is actually proving (in the mathematical sense) that the logical properties are consistent over the whole program or library. Removing type checking (through weak or dynamic typing) is equivalent to trusting the programmer for checking these logical properties themselves. This means trusting a *homo sapiens*, for being consistently and carefully logical all the time; which is absurd and very ill-advised [21]. We see expensive consequences of this irresponsible behavior for instance when the security community exhibits exploitable software flaws, recent findings are even brought to the attention of bioinformaticians [22]: while the article suffers from sensationalism, it highlights the need to be sure that any piece of software behaves correctly for all possible inputs, including maliciously crafted ones. Software testing, no matter how thorough, only gives confidence on tiny subsets of possible program executions. Of course, while the software safety research community has made immense advances, as of 2017, fully formalized and proved software is still out of reach for large-scale, open-source, and under-funded software engineering. Testing remains important, but it has to be in addition to static and “as strong as possible” typing.

Given the above considerations, actual choices for implementation were limited, but OCaml stood out. It is equipped with a very advanced but *practical* type-system [20], [23]. The system was carefully designed with speed *and* correctness in mind and offers a lot of flexibility. The community includes very successful industrial users for whom safety and security matter [24], [25], and has contributed great tooling. OCaml is also more future-proof; as the next generation of safer programming languages and tools, like Coq [26] or F* [27], are often written and/or extract code to OCaml.

### Next In This Paper

We are now going to see deeper into the proposed systems, the remaining of this paper is organized as follows. We start at a lower-level (i.e. close to the computing environment) with Ketrew and Coclobas, our infrastructure for workflow automation, in section. Then, we climb the abstraction hierarchy through Biokepi, our library for building bioinformatics pipelines, up-to it’s EDSL based on recent computer-science research (section). In section, we quickly present an open-source and quite feature-rich use case: Epidisco. We finish with related (section) and future (section) work.

## Ketrew and Lower-Level Considerations

*Ketrew* stands for “Keep Track of Experimental Workflows,” we detail first its low-level design and implementation, and then its pipeline-construction API. We end this section with *Coclobas* which stands for “Configurable Cloudy Batch Scheduler.”

### The Ketrew System

The initial design and first versions of Ketrew had two modes of operation: a “client-server” — default/normal — mode, and a “standalone” one. The standalone mode has been dropped since; it was meant to allow users to quickly setup and try running workflows by not having any server or daemon (Ketrew itself or a database) in place, and doing everything with a single Unix process. This *standalone* behavior exists in other workflow automation tools (it is the default for utilities similar to make or bigger systems like Luigi [28]). We dropped its support for various reasons, among them: i) the interactive web-based user interface (a.k.a. “WebUI”) has proven very useful especially for beginners and it can only be used when a server is running; ii) the operation of the standalone mode was confusing users regardig what they can or cannot do “concurrently” (it was based on the *Sqlite 3* database which does not fully support inter-process concurrency [29]). Instead we have insisting on making the client-server mode as easy and flexible to setup as possible [30].

The Ketrew application is hence one simple Unix binary which contains the service logic (called “the engine”), a web-server, a command line interface (CLI), packed together with the Javascript code (compiled from OCaml thanks to `js_of_ocaml` [31]) and the style-sheet (CSS) of the WebUI. The system can be installed and used on any Linux system or on Mac OSX and, if needed, everything can be done as a regular Unix user. The server-side component is designed so that the application can be abruptly stopped or become offline while workflows are running, and restarted seamlessly in the right “state.” The HTTP API and the WebUI can be served over plain HTTP or with TLS (both the classical OpenSSL library, and the safer nqsb-TLS [32] can be used). There is also a semi-interactive text-based user interface which communicates with the web-server using the same protocol as the WebUI; users usually prefer the latter but the TextUI is for example useful when institutional firewalls or virtual-private-networks (VPN) get on the way of regular web-browsing.

Usually, scientific computing clusters run a system component to abstract the setup and monitoring of programs for regular users; this is known as a *batch/job scheduler*. A workflow engine needs to interface with these systems to actually run commands and scripts on the targeted infrastructure. Ketrew adopts a plugin architecture for the implementation of the communication with these system called “backends.” Any user can write new backends and load them with the system (and if deployment of dynamically loaded modules is a problem; one can easily create a Ketrew application binary statically linked their plugins). We ship various backends: Platform LSF [33], PBS/Torque [34], YARN [35] (with which we can run both Hadoop applications like Spark applications or regular shell-based jobs), and a “daemonization” plugin capable using two methods (one for more “standard” Unix hosts based on the nohup and setsid programs, and one based on generated Python scripts — for Mac OSX hosts).

At a pipeline-level, the choice of backend is attached to lowest-level jobs; each step can choose the plugin it uses. Unlike many other workflow-engines where the backend is set at a global level, cross-infrastructure workflows are hence easy to write and setup (e.g. replicating data from one cluster to another, or running some steps outside of the scheduler for speed-up).

Ketrew’s engine (and its plugins) can communicate with the infrastructure directly on the current host (as simple system calls) or over SSH connections. Indeed, users can setup password-less SSH access in order to, for instance, run Ketrew on their laptop and manage workflows on one or more university clusters, even when they cannot have a long-running server on their institution’s infrastructure.

As a *failed* experiment, we also built with a system for users to setup so-called “control master” reusable SSH connections from a shared Ketrew server. Even though OpenSSH is specifically designed to make it hard for users to “script” the client, we managed to get a prototype working: a Web interface to setup SSH connections (working even with password-only or 2-factor authentication schemes). We then hit multiple unexpected failures from OpenSSH, it proved not reliable enough for heavy duty use of the system (there are hard limitations on connection multiplexing [36]).

### Ketrew’s EDSL

We provide a very flexible EDSL-based API to construct workflows. The EDSL is provided as a simple and pure OCaml library. All the constructs of the language are used to build an immutable graph data-structure. This helps the users organize their (partial) workflows in a modular way and with their own domain-specific abstractions. The resulting functions and libraries are then nicely composable and easier to reason about.

The submission of the workflow to the server is done with one simple function that serializes the resulting data-structure to a JSON object [37] and sends it to the server over HTTP(S). The Ketrew engine performs a node equivalence search before starting any jobs; this means that unless explicitly disabled, nodes that attempt to produce the same result will be “merged.”

On a semantic level, the API of the EDSL is designed to help the user build a graph with three kinds of edges. The nodes contain most of the information (how to run the step, how to check whether it is already done or successful, and much more meta-data) and the edges are either *dependencies* (the most common way of constructing workflows) or mechanisms to react to success or failure of the nodes (for instance, we can define “clean-up” workflows that are activated when a step fails).

Workflow nodes are meant to ensure logical “conditions.” These are expressed thanks to a (much smaller) EDSL of boolean expressions whose base terms are checks on precise system conditions (e.g. that a given file or a file-tree structure exists, or that a shell-command returns a given code, etc.). As we started using the first iteration of the EDSL for larger scale pipelines, the importance of the object(s) of these condition-expressions proved to be bigger than anticipated. Hence, in the second (and current) major version of the EDSL, we use a type-parameter to pass the products of the nodes around the programs in a strongly typed way (cf. Parametric Polymorphism [38]). A “product” is then an abstraction of what a workflow node ensures. Using different type-parameters for different kinds of workflow-nodes helps us make senseless code impossible to express; those constructs are extensible by user-code. For instance, within Biokepi, we cannot use a node which produces Bam files in a place where we expect VCF files; those products are defined in the library as they are bioinformatics-specific. Differently typed nodes, can still be be packed together, e.g. as a list of dependencies, thanks to existential types [39]; they become “edges” in the pipeline graph.

### The Coclobas Backend

In practice, our team has run Ketrew workflows first on a Plaform LSF cluster, and then on a YARN-based Hadoop cluster, until moving to Google Cloud’s infrastructure. We consider relevant and interesting to report on this experience.

We first were guided by the goal of utilizing the infrastructure as fast as possible. Hence we wanted to quickly set-up/update/destroy familiar PBS/Torque [34] clusters with shared file-systems. As we tried and were disapointed with the reliability of existing solutions (e.g. Elasticluster [40]), we decided to use Ketrew workflows for the task (process often known as “dogfooding,” i.e. using our own product to stress-test it). The resulting project was called Stratocumulus [41]; configurable workflows which could set up shared computing infrastructure on the Google Compute Engine.

The above project got us and our users to very quickly get started leveraging the infrastructure but it was not cost efficient. Indeed, building “classical” computing clusters on cloud infrastructure most often means keeping compute nodes up and running even when there is no work to do. Manual update or destruction from users when they are done cannot always be counted on; for instance, we observe that when a workflow finishes on Friday night, users will not collect their results and clean-up their resources until Monday morning. This lead us to investigate ways of using auto-scaling capabilities provided by some Google Cloud components: we created a new job scheduler, Coclobas, accepting jobs over HTTP and scheduling them on elastic Kubernetes [13] clusters (as deployed by the *Google Container Engine* — GKE). Later, the project was extended to schedule container-based jobs within *AWS-Batch* and on local Docker installations. Note, that while the main client is the Ketrew plugin, one can submit jobs to Coclobas without Ketrew. Coclobas also takes care of working around various idiosyncrasies of Kubernetes: it keeps track of the logs (which Kubernetes can easily loose), and, it throttles submissions and retires after failures to limit the impact of overloading of the Kubernetes server. The API also simplifies like the use of the “secrets” feature to pass custom information to containers (e.g. a script to run), or the setup of arbitrary NFS mounts. Coclobas can use an existing server or manage a fresh one using the Google Cloud client.

## Abstractions in Biokepi

Ketrew and Coclobas are also used for workloads not related to bioinformatics (like system administration [41], building documentation, etc.) but the main strength of Ketrew’s API is being an embedded DSL in a powerful language like OCaml. This allows users to build modular abstractions that fit their application domain using proper software engineering. Our abstractions for bioinformatics, i.e. the Biokepi library, are detailed in this section.

Bioinformatics workflows in Biokepi are organized in two layers: the lower-level layer consists in a catalog of bioinformatics tools wrapped as Ketrew workflow nodes. The second layer is the higher-level `Pipeline_edsl` module; it is an embedded language to write workflows very concisely with help from precise types.

### The “Tools” API

At this lower-level, we use already proper types to give stronger semantics to the tools’ parameters, and add constraints and invariants. We have for instance abstractions of FASTQ [42] single-end or paired-end sets of files, or Bams [8] recording their reference genome their sorting status (coordinate, read name, etc.). We also use these modules to encode our slowly acquired knowledge about the idiosyncrasies of biomedical software. Having a proper programming language also simplifies the implementation of decently complex performance improvements like automatically generating *“scatter-gather”* parallelizations of some computations or replacing partial of workflows with “chained” shell pipes.

The workflow nodes are designed to be able to make everything *“restartable”* and leverage Ketrew’s semantics to share intermediate results as much as possible. All of this is built around the “Machine” abstraction, a module defining the computing infrastructure and environment (bioinformatics software and data) to simplify the implementation of very portable and configurable workflows. This environment can take care of most software installations, but it is easily configurable; for example one can leverage software already present on the user’s infrastructure. Similarly one uses the abstraction to configure the access to “reference-data,” by default through downloads from public sources.

### The Typed-Tagless Final Interpreter

The second layer is the higher-level `Pipeline_edsl` module; it is an embedded language to write workflows very concisely with help from precise types. The EDSL hides out workflow steps that we consider “boilerplate” (like indexing and sorting Bam files, preprocessing reference genomes, locally installing software, etc.).

The first version of the EDSL that we implemented was based on a generalized algebraic data type (or GADTs, [43]). The module (that we maintain for backwards compatibility for now) proved very practical for end users as long as they did not want to extend the language without modifying Biokepi itself. Moreover the GADT-based implementation lacked modularity and could have grown to proportions difficult to maintain. Therefore we decided to rewrite this component to be based on recent research on *typed tagless-final interpreters* [18], [44]. While being slightly more complicated to approach the new implementation provides extensibility while being well-typed. Examples of extensions are detailed in section.

The pipelines written using Biokepi’s high-level EDSL can be compiled to various backends. Of course the main compilation target are Ketrew worklow nodes (using the lower-level modules of Biokepi), but we can also generate high-level graph descriptions (i.e. at the level of bioinformatic-tools/semantics) using the “dot” language from the Graphviz project (cf. example in figure 2). There is also a compiler to JSON files allowing implementation-independent traceability (and potential reproducibility) of the workflows (human-and-computer-readable exact descriptions of the whole pipeline). Biokepi still allows to “break” the abstraction barrier and write workflows with lower-level functions manipulating Ketrew workflow-nodes through extensions of the EDSL and users can write their own compilers/interpreters.

Because of our main application domain, the tools avaialable in Biokepi are for now mostly focused on cancer genomic pipelines but we welcome contributions from any sub-field of bioinformatics. The following section presents Epidisco a configurable pipeline based on Biokepi’s EDSL.

## Use Case of Epidisco and the PGV Trial

In the context of our participation in a Personalized Genomic Vaccine clinical trial (NCT02721043 [16]), we have developed Epidisco, a family of pipelines for selecting vaccine peptides targeting cancer mutations. Epidisco is a Biokepi pipeline that produces ranked peptides from the outputs of the sequencer (by default FASTQ data, but we can also start from BAMs and realign them automatically); it is designed to run with an arbitrary number of normal, tumor, and tumor-RNA samples, plus optional HLA-typing information (if not provided the pipeline computes it) [16], [17], see figure 1. The pipeline produces an HTML report that we can serve to our collaborators together with the results.

**Figure 1:**
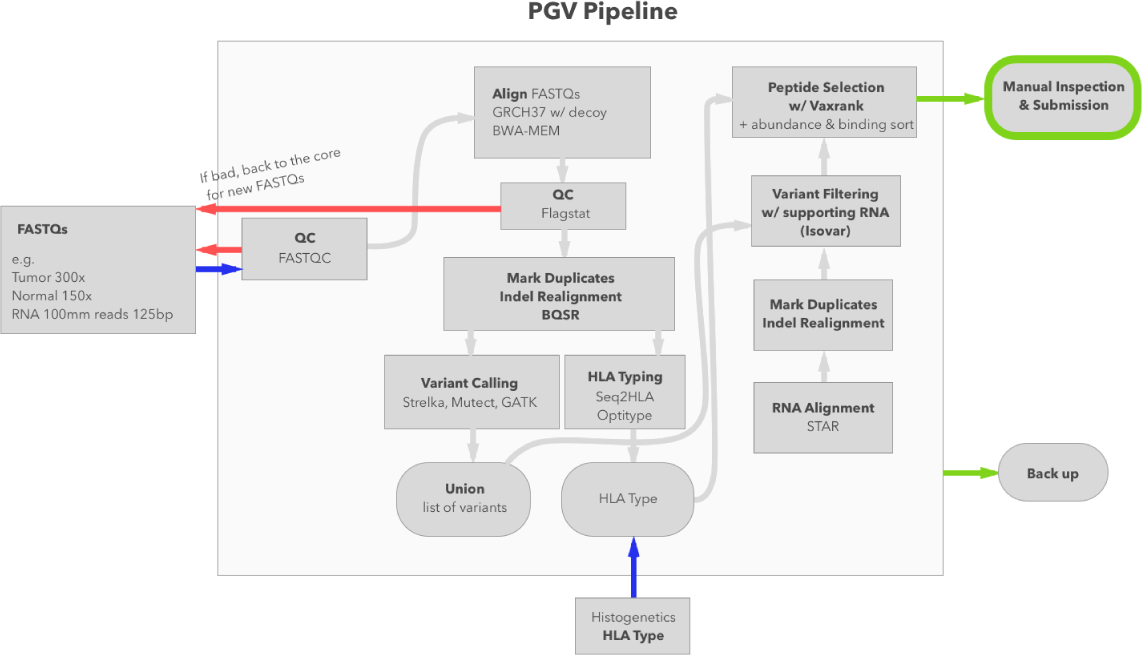
The PGV/Epidisco Pipeline. High-level diagram of the pipeline that we currently have running in production.

**Figure 2:**
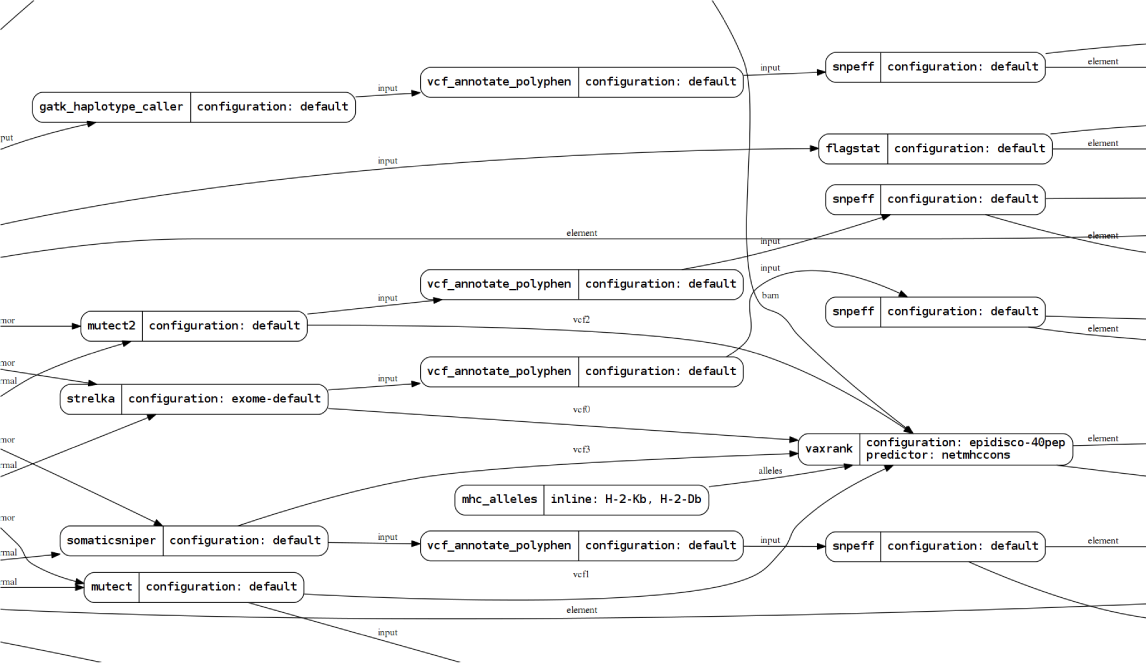
Biokepi-EDSL Example Pipeline. Excerpt from the Graphviz-based rendering of a pipeline which uses Biokepi’s EDSL.

Epidisco fully utilizes the flexibility provided by the Biokepi EDSL. We extend it with custom constructs that are specific to the particular application (like the “saving” of intermediate results or construction of the final report web-page). The big and growing amount of options that can modify the pipeline enables us to maintain a “production” pipeline while allowing various experimentations; the high-level EDSL makes the code still reasonably readable and easy to understand. Moreover using proper software engineering at the pipeline-level makes perilous changes much easier; for instance, one can see that the logic used to experiment-on and implement the fix in the pull-request `#119` [45] would have been much harder to express in a custom workflow DSL or with a weak programming language.

## Related Work

The literature and the open-source world contain many workflow engines and computational pipeline tools. Some of them are biology-specific specific and some are more generic. Before starting working on this family of projects we reviewed and tried a few (including commercial software, although inadmissible for security and long-term dependence reasons). None were fully satisfying although we borrowed ideas and lessons from many of them. Note that Spjuth et al. previously published a biased but relatively thorough review paper on the matter [46].

Given the diversity of the analyses and of the software environments, most workflow management tools that aim at specializing (in bioinformatics or other) end up not being flexible enough and having to implement many edges cases one by one. A good example is the famous Galaxy [47], which is quite inflexible while being very thorough has required many human-years of effort to be implemented and by the time we tested still didn’t present the reliability and flexibility that a small fast-paced team requires. QuickNGS [48] is actually a LIMS (Laboratory Information Management System) that happens to implement a simple workflow engine to run a predefined set of tools; this can be useful for core facilities that follow the same overall functioning. ExScalibur [49] is a set of automated pipelines for whole exome data analysis, that is implemented in the custom DSL “BigDataScript” with very little abstraction power. COSMOS [50] at least uses an EDSL, within Python, but its model restricted to map-reduce-like workloads (direct acyclic graphs of shell commands producing files). Azkaban [51] is a heavy-weight workflow manager used for *“several years at LinkedIn.”* Its design is specialized for quick-running Hadoop pipelines although extensible through Java plugins; workflows are defined very awkwardly as key-value configuration files. Still extremely specialized, Makeflow [52] is actually a set of tools which provide abstractions for particular computational patterns (e.g. “Map-Reduce” is such a kind of abstraction). GXP Make [53] (based on “GXP Shell” [54]) is a fun (ab)use of GNU-make as it provides a shell mksh that intercepts make’s calls to run distributed workflows. Biomake [55] is another extension of the venerable make tool by making it more programmable thanks a Prolog execution engine. Swift [56] is a C-styled custom, hence limited, language used for encoding Makefile-like dependency graphs which can be run on various platforms. Taverna [57], now an Apache Incubator project, attempt to be a “Graphical Programming Environment” to define workflows and share them through MyExperiment.org project.

Other tools, have embraced the “Embedded Domain Specific Language” bandwagon but most often fell short on the reliability and expressivity aspects. For instance, BPipe [58] while claiming no need for programming experience, is also an awkward EDSL within Groovy, an unacceptable Python-like language for the Java Virtual Machine. Snakemake [59], Ruffus [60], and Luigi [28], are Python EDSLs, the latest being most advanced one, while quite Hadoop-centric. Similarly, Pwrake [61] is an extension of Rake (Ruby EDSL-ish build system) to run “builds” in parallel. Queue (part of the GATK [62]) on the other hand is a Scala-based library; the extreme object-orientation brings the verbosity of Java while not trying to improve on the type safety.

## Future Work

Like a almost any open-source family of projects, more future work can be envisioned than humanly achievable.

One the lower-level aspects we are actively working on extending the catalog of backends that Ketrew can utilize: after the Google Container Engine and the “Local Docker” setup, we are now improving support for Amazon AWS.

For the workflow engine itself, after a few recent performance and scalability improvements, the main point we want to improve is now the WebUI. We want to make it extremely easy to build custom “job submission interfaces” from a high-level and embed them in Ketrew’s WebUI. Multi-scale graphical visualization of large workflow graphs is also both a very appealing feature and an interesting, surprisingly “open,” problem to work on.

Ketrew’s API for writing workflows has proved to be very practical and scalable but the actual shells commands run by workflow-nodes are still mostly untyped strings, the module provides a few higher-level constructs but we want more “typed programming” abilities. This is why we have been developing Genspio [63], a *typed* EDSL to generate POSIX shells scripts; we avoid the shell’s “escaping hell” and provide a more composable API. Genspio has been recently released, and stress-tested in a systems-administration context; the next step being its integration into Ketrew.

At the level of Biokepi, in addition to the expansion of the “tool catalog,” it may be interesting to add more *formal* information into the types of the constructs of the EDSL (e.g. whole genome or exome, species, RNA/DNA). Of course, we also want to explore, at any level of the system, the progressive introduction of more precise and stronger formal guarantees through dependently typed approaches [26].

